# genomicSimulation: fast R functions for stochastic simulation of breeding programs

**DOI:** 10.1101/2021.12.12.472291

**Authors:** Kira Villiers, Eric Dinglasan, Ben J. Hayes, Kai P. Voss-Fels

**Affiliations:** Queensland Alliance for Agriculture and Food Innovation, The University of Queensland, St Lucia, QLD, Australia; Department of Grapevine Breeding, Hochschule Geisenheim University, Geisenheim, Germany

**Keywords:** breeding program simulation, meiosis simulation, breeding program design, genomic selection, R package, C language

## Abstract

Simulation tools are key to designing and optimising breeding programs that are multi-year, high-effort endeavours. Tools that operate on real genotypes and integrate easily with other analysis software are needed to guide users to crossing decisions that best balance genetic gains and diversity to maintain gains in the future. This paper presents genomicSimulation, a fast and flexible tool for the stochastic simulation of crossing and selection on real genotypes. It is fully written in C for high execution speeds, has minimal dependencies, and is available as an R package for integration with R’s broad range of analysis and visualisation tools. Comparisons of a simulated recreation of a breeding program to the real data shows that the tool’s simulated offspring correctly show key population features, such as linkage disequilibrium patterns and genomic relationships. Both versions of genomicSimulation are freely available on GitHub: The R package version at https://github.com/vllrs/genomicSimulation/ and the C library version at https://github.com/vllrs/genomicSimulationC/

---

When real breeding schemes and genetic improvement programs are high-effort, multi-year undertakings, simulation tools offer valuable opportunity to judge feasibility or balance trade-offs in design so as to get the most out of the real programs (Li et al., 2012). The available genetic simulation tools as described by Sun et al. (2011) have tended to be standalone tools with limited opportunities to incorporate external analysis tools, and/or tools that generated artificial genomes from parameters rather than using real data. While this is helpful for statistical analyses or broad evaluations of breeding strategies (for example, Hickey et al. (2014)), one of the key problems in optimising breeding scheme design is parent selection, which is highly dependent on the actual parents that are available to the breeding program (Sun et al., 2011; Li et al., 2012; Bernardo, 2020). Bernardo (2020) makes a strong case for the need for simulation tools focused on operating on real genotypes data, real genome maps, and real phenotypic performance data.

This paper introduces genomicSimulation, a fast, flexible R package for the stochastic simulation of breeding programs on real genotype data. It was built to provide high-speed crossing simulation capabilities that can be easily integrated with R’s wide range of visualisation and genetic analysis tools. genomicSimulation loads and runs based on user-provided marker maps, founder genotypes, and (optional) lists of allele effect values, so can easily simulate breeding schemes on real maps and candidate founders. It simulates meiosis without mutation on alleles at discrete positions provided in the genome map. The linkage phase of resulting genotypes is tracked. genomicSimulation works as a scripting tool, with functions for performing targeted crosses, random crosses, doubled haploids and selfing. genomicSimulation’s inbuilt genotypic value calculator uses an additive model of marker effects. Alternatively, users can script their own custom selection methods in R. The package has no dependencies beyond C standard libraries. All core functionality is written in C in order to achieve high execution speeds.

For even faster simulations, genomicSimulation’s underlying C library (in itself a fully-functional simulation tool) is also available. It is distinguished from the R package only by the lack of default parameters and lack of R vector data output options.

The package was originally developed for use on self-pollinated crop species, but its flexible set of crossing operations allow it to be used more broadly in outcrossing species. genomicSimulation has full documentation, user guides, and is in ongoing development.

## Methods

genomicSimulation provides a toolset of ‘building block’ scripting functions. Users have full flexibility to intersperse requests to perform a cross (and produce new offspring) with commands to restructure groups (to perform selection or restructure the breeding pool).

The first step in using genomicSimulation is a call to the setup function,load.data(). Two data files are required to set up the tool: a matrix file contain the alleles of at least one founder genotype at a set of discrete positions, and a linkage map to situate those positions in the model of the genome. If the phase at heterozygous positions in the founder genotypes is not known, it is randomised on load. Alleles can be any non-space character, which allows for the package to be used on defined and known Mendelian genes as well as SNPs.

The simulation stores every simulated genotype as a sequence of characters in memory.

To create a cross, the simulation generates a gamete independently from each of the two parents. No distinction is made between male and female parent. Karlin and Liberman (1979)’s count-location method is used to simulate meiosis. Specifically, the following process is followed (Figure 1). First, the number of crossovers to occur in each chromosome is drawn from a Poisson distribution whose parameter is the distance in Morgans between the first and last positions tracked on that chromosome (Figure 1a). The positions of those crossovers along the chromosome are then drawn from a uniform distribution (Figure 1b). Finally, a random logical value (0 or 1) is drawn to choose which of the two possible gametes to use (Figure 1c).

The count-location method of simulating meiosis ignores crossover interference. The specific distributions chosen add the additional assumptions that the number of crossovers is proportional to the chromosome length, and that crossovers are uniformly distributed along the chromosome. This choice of distributions and assumptions is shared with existing simulation tool Plabsoft (Maurer et al., 2007). Functionality to predetermine crossover sites or probabilities, if for example linkage disequilibrium block structure is known, may be added in future.

During setup, there is also the option to load an input file containing the numeric effects of particular alleles. This enables the internal breeding value calculator. Breeding values are calculated using an additive model of effects: the breeding value of a given genotype is the sum across all marker positions of each marker’s two alleles’ effect values (if such allele effects are loaded). This function can be used to compare different breeding schemes with regards to rates of genetic gain for a defined trait, or to assess the genetic merit of specific crosses for the traits for which they have estimated marker effects, as a basis for crossing decisions.

### Features

At any time after the initial setup command, more external genotypes can be loaded using the command load.more.genotypes(), and the simulation’s optional stored set of marker effects can be substituted for another with load.different.effects(). With these features, users can simulate the introduction of new genetic material into an ongoing breeding program, calculate genotypic values for multiple traits with separate sets of marker effects, and/or manually simulate environmental fluctuations across years and/or locations by using different sets of marker effects.

A range of functions are available for simulating the production of offspring. These include cross.randomly(), self.n.times(), make.doubled.haploids(), and cross.all.pairs(). To carry out specific crosses or crossing plans, users can call cross.combinations() with vectors laying out the parents to cross.

Every genotype loaded or produced in genomicSimulation is allocated to a group. Every function in the R package returns 0 if it does not create or modify any groups, or the group identifier (a positive integer) of whatever group was populated. Mixing and separating groups allows for significant flexibility in regards to simulating multi-generational breeding pools, or having several interacting streams in the breeding program. Inbuilt (non-custom) options for group manipulation include combine.groups(), break.group.into.families(), break.group.into.halfsib.families(), and a range of functions to randomly partition a group (which can be used to demarcate male/female offspring, or to create sub-groups of specific sizes for complex breeding program designs), alongside custom group manipulation using make.group().

The function select.by.gebv() performs truncation selection on breeding values calculated by the internal breeding value calculator. An interface for custom selection methods is also offered: this involves using data from see.group.data() and some R scripting to identify the best individuals, followed by passing those individuals’ indexes to make.group() to move them into a new ‘selected’ group.

Groups and genotypes persist until explicitly destroyed. Therefore, mixed-generation crossing operations are possible, and users have control over their memory resource usage.

Simulated data can be saved to tab-separated text files or pulled into the user’s R environment as vectors. The current data types available as output are genotypes, haplotypes, allele counts, immediate parents, full known ancestry, breeding values (calculated using the internal additive trait), and local breeding values over certain blocks/subsets of positions (calculated using the internal additive trait effects).

More in-depth description of features and usage examples are provided in the R package vignette, R documentation, C library guides, and C library documentation. These can be downloaded with the package or accessed at the package GitHub links in the abstract.

### Run Speed

Table 1 shows some average runtimes of simulation tasks. Most constraints on speed are imposed by file read/write operations, so scripts that minimise the number of outputs saved will run faster.

**Table 1:**
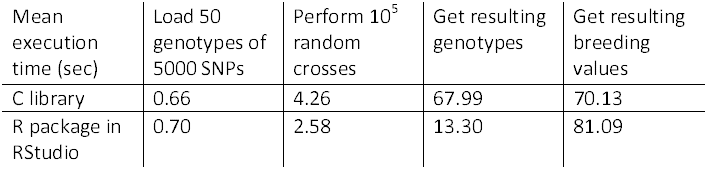
Average execution time (in seconds) across 6 repeats of tasks in genomicSimulation v0.2, running on a consumer-model laptop with 8GB RAM and Intel i5-7200U CPU @ 2.50GHz. The tasks timed are: (1) loading 50 genotypes of 5000 SNPs, (2) perform 10^5^ random crosses between those genotypes with one offspring each, (3) save the 10^5^ genotypes from task 2 to a file (for the C version) or an R dataframe (for the R version), and (4) calculate then save the breeding values of the 10^5^ genotypes from task 2 to a file (for the C version) or an R dataframe (for the R version). Note that the R version of the tool also shares the C library’s functionality for saving output to files. The time taken to save output to files is comparable across R and C versions.

| Mean execution time (sec) | Load 50 genotypes of 5000 SNPs | Perform $10^5$ random crosses | Get resulting genotypes | Get resulting breeding values |
| --- | --- | --- | --- | --- |
| C library | 0.66 | 4.26 | 67.99 | 70.13 |
| R package in RStudio | 0.70 | 2.58 | 13.30 | 81.09 |

## Results

### Example Simulation

To provide an insight into the effectiveness and flexibility of the genomicSimulation tool, a sample script for simulating a simple wheat breeding program with non-overlapping generations is shown in Box 1. The simulation was initiated with genotype data for 50 real founder lines (Supplementary Figure S1), and the effect values initialised with values calculated from phenotypic data for yield.

The simulated breeding program was divided into cycles. The first stage of each cycle involved random crosses between founder inbred lines (using cross.randomly()) to generate new recombinants. One selfed progeny from each recombinant was ‘grown’, and phenotyped at 10% broad-sense heritability for the yield trait. The top 20% of the generation by phenotype was selected to pass on to the next generation. One selfed progeny of each of these was ‘grown’ and phenotyped at 40% heritability for the yield trait. The top 50% of that generation were selected and progressed by selfing to the last generation of the cycle. These final progeny then served as the founders of the next cycle. Figure 2a includes a visualisation of the steps of the simulated breeding cycle.

**Figure 2:**
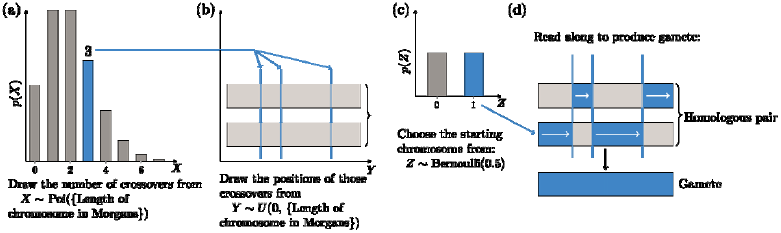
Meiosis simulation procedure in genomicSimulation uses a count-location strategy. The following steps are performed for every homologous pair of chromosomes. (Blue coloration shows a sample procedure.) (a) First, the number of crossovers is drawn from a Poisson distribution with expectation matching the length of the chromosome in Morgans. (b) Next, the positions of each of those crossovers are drawn uniformly across the length of the chromosome. (c) A final random draw determines which of the two resultant gametes is chosen. (d) The gamete is created by reading along the chosen starting chromosome, swapping to the other of the pair whenever a crossover point is encountered.

In this simulation, genomicSimulation’s groups were used to represent the new population of plants grown each generation, and also to pull the selected plants out of the generation’s wider generated population.

Two conditions were tested in simulation – one where selection was across the whole set of lines in that generation, and one where selection was performed within each family (the set of seeds sharing the same parent crossed plants).

The phenotype on which selection was performed was simulated in a user-scripted R function, to demonstrate the mechanism of designing custom selection methods in genomicSimulation. Phenotype was simulated as *P* = *G* + *E*, where the environmental contribution *E* is a normally distributed variable with mean 0 and variance *V*_*e*_. Given a broad-sense heritability value for the particular selection stage, the variance *V*_*e*_ can be calculated by rearranging:

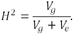

The results of running the script were consistent with expectations. Figure 2b shows that the genetic value of the yield score in the population increased with each cycle, because of the correlation between the phenotype and the heritable genetic trait. The condition where selection was performed across all plants showed a higher rate of gain than the condition where selection was within families, because it could select more plants from good families and therefore increase the proportion of good alleles in its population faster. Figure 2c shows the variance in genetic value scores decreased as cycles increased, as the proportion of beneficial alleles in the population and the proportion of plants with many of these alleles increased. Selecting within families kept this diversity measure higher than selecting across the whole population.

**Box 1:**
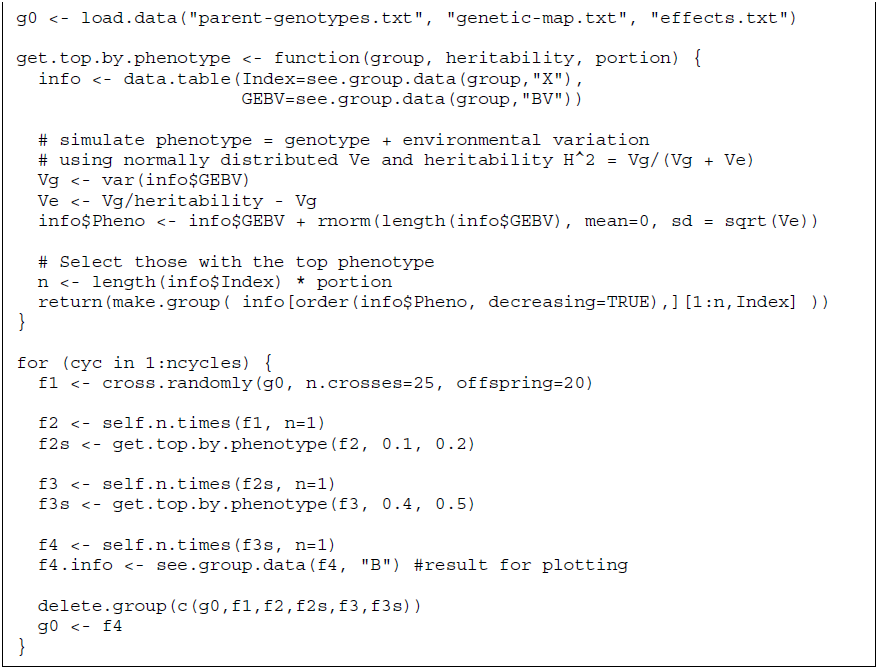

*R script to simulate a simple breeding program using genomicSimulation. The program uses non-overlapping cycles and selects after the first two selfing steps with different accuracies. This version implements the “across all lines” selection condition results shown in Figure 2b-c; the script for the “within families” condition calls break.group.into.families() after generating the F1 crosses, then runs the rest of the script commands independently for each family group produced by the break.group command.*

### Validation

A structured durum wheat population from Alahmad et al. (2019) was simulated using genomicSimulation to assess the tool’s ability to recover the genetic structure found in the empirical data set. The structured population was developed through the nested association mapping (NAM) design presented in Figure 3a. This design was simulated with genomicSimulation using the cross.combinations() and self.n.times() commands to produce 10 families of 100 genotypes in the final generation.

**Figure 3:**
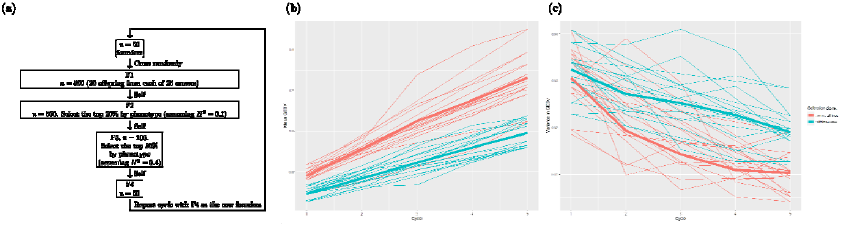
(a) Diagram of each cycle of a simple breeding program plan (see Box 1 for the genomicSimulation implementation). The program was simulated using genomicSimulation. (b) Mean and (c) variance in the population’s genetic breeding value are shown for each replication (thin lines) and averaged across replications (thick lines). In the first condition, the best phenotypes across the entire population in the relevant generation are selected, while in the other condition, the best phenotypes in each family are selected.

**Figure 4:**
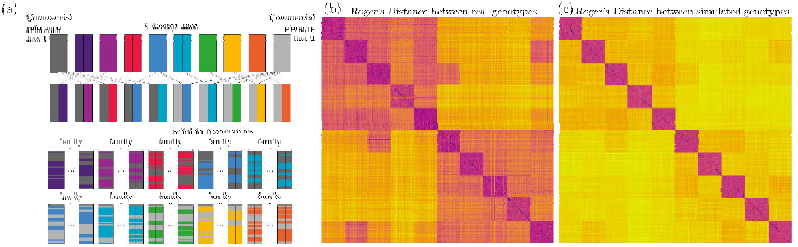
(a) The crossing plan of a structured population developed by Alahmad et al. (2019). The matrices of Roger’s Genetic Distance between all (b) real imputed genotypes, (c) simulated genotypes, of the final-generation offspring resulting from that crossing design.

R packages SelectionTools v. 19.4 and ComplexHeatmap v. 2.9.3 were used to produce the heatmaps of genetic distance shown in Figure 3b-c. The population structure of ten family relationships and two shared-elite-parent relationships is clearly visible in both simulated and real genotype heatmaps. The distributions of genetic distances also show the same profiles (Supplementary Figure S2). However, the real population shows higher variation within families, as shown in the wider range of base colours in the heatmaps and in Supplementary Figure S3. The real population was developed with selection for several traits during line development, including height variation, reasonable flowering time and early vigour (S Alahmad, personal communication, 27 January 2021). These selection pressures were not mimicked in this simulation, and so may well account for the differences in level of variance observed.

## Discussion

Simulation is a valuable tool to investigate the choices needed to carry out a breeding program. Various simulation tools in this area exist. Some, such as QU-GENE (Podlich and Cooper, 1998) and ADAM-Plant (Liu et al., 2019), are standalone programs, but an increasing number are R packages. These include Plabsoft (Maurer et al., 2007), BSL (Yabe et al., 2017), MoBPS (Pook et al., 2020), and AlphaSimR (Gaynor et al., 2020). genomicSimulation joins these ranks as a simple, flexible, freely-available tool designed for simulating from real genotypes and genome maps at high speeds. It is available as a standalone command-line tool or as a set of R scripting commands.

Its approachable syntax, easy interoperability with other R libraries, and automatically-fast execution (thanks to the underlying C library implementation) make it a useful tool to simulate, test, and compare breeding strategies. At present, it offers an additive internal breeding value calculator, although this offering will be expanded to include non-additive genetic effects. It is expected that users script more complex genetic evaluation methods themselves, for example, scripting a simulated phenotype (as in the first example in this article), or recalculating SNP effect values (using one of R’s specialised genomic selection packages) and substituting them into genomicSimulation using load.different.effects() at relevant points during the breeding program simulation.

R’s popularity in biological data analysis mean it houses a range of cutting-edge genetic analysis and genomic selection packages with which users may already be familiar. Developing a breeding program simulation in base R, however, requires significant effort and planning, and may produce a slow-running tool. genomicSimulation provides an engine to carry out simulations according to breeding programs that can be scripted in R. A key advantage of genomicSimulation is that is uses real founder genotypes, genetic maps and incorporates other biological features of target species, which adds to the immediate usability of simulation results.

genomicSimulationshould be straightforward to install thanks to its lack of dependencies beyond C standard libraries. Documentation and use guides are available. Development is ongoing.

## Acknowledgements

The authors thank Samir Alahmad and Lee Hickey for providing access to the wheat NAM population dataset.

## Data Availability

Code and installations of both versions of genomicSimulation are freely available on GitHub: The R package version at https://github.com/vllrs/genomicSimulation/ and the C library version at https://github.com/vllrs/genomicSimulationC/. The scripts and founder genotypes used in the example simulations in this article are available upon request.

