## Supplementary Figures for "genomicSimulation: fast R functions for stochastic simulation of breeding programs"

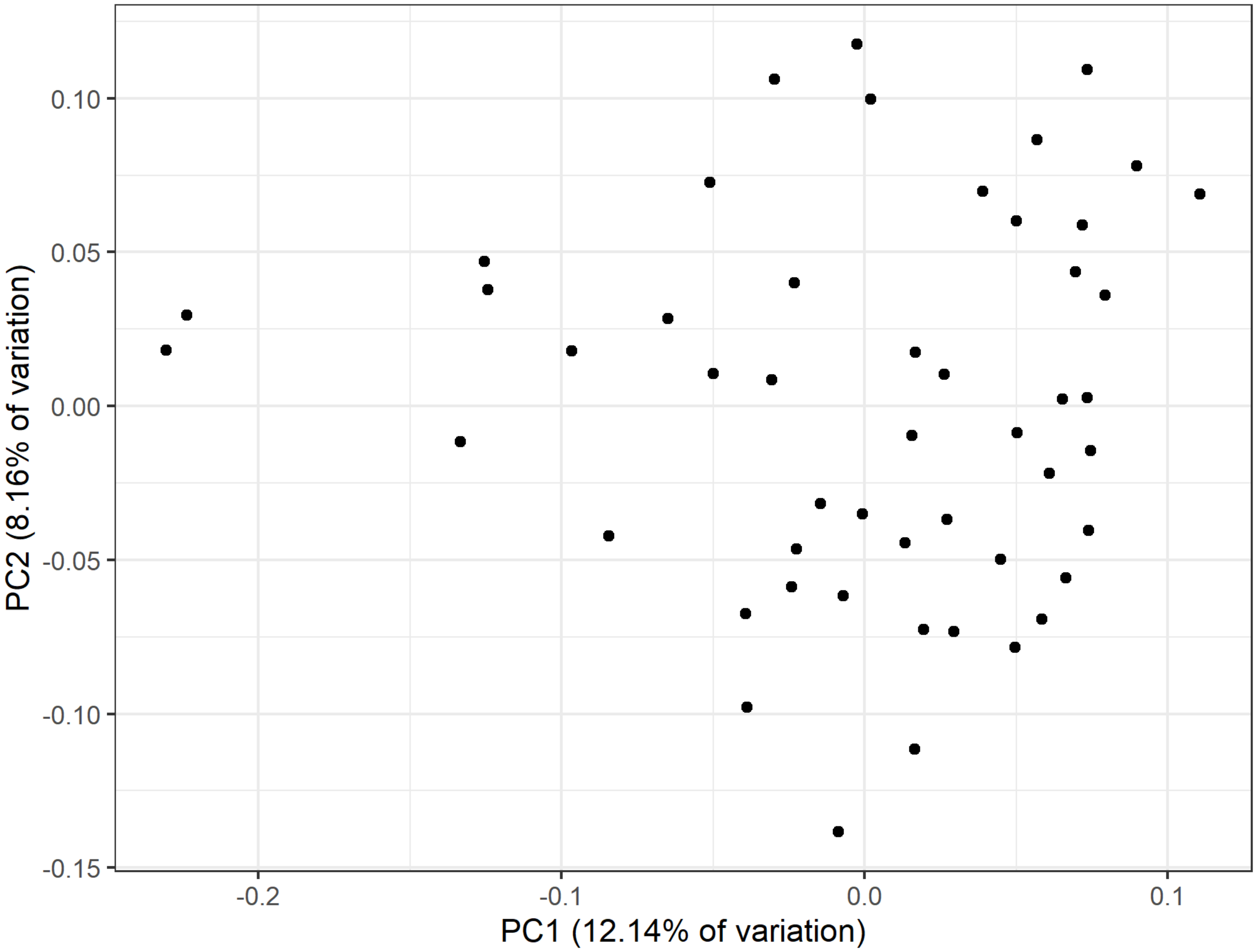


Figure S1: Principal component plot of the 50 wheat genotypes used as founders in the example simulation (Figure 2 in article). These founders were a set chosen for encompassing a wide range of diversity.

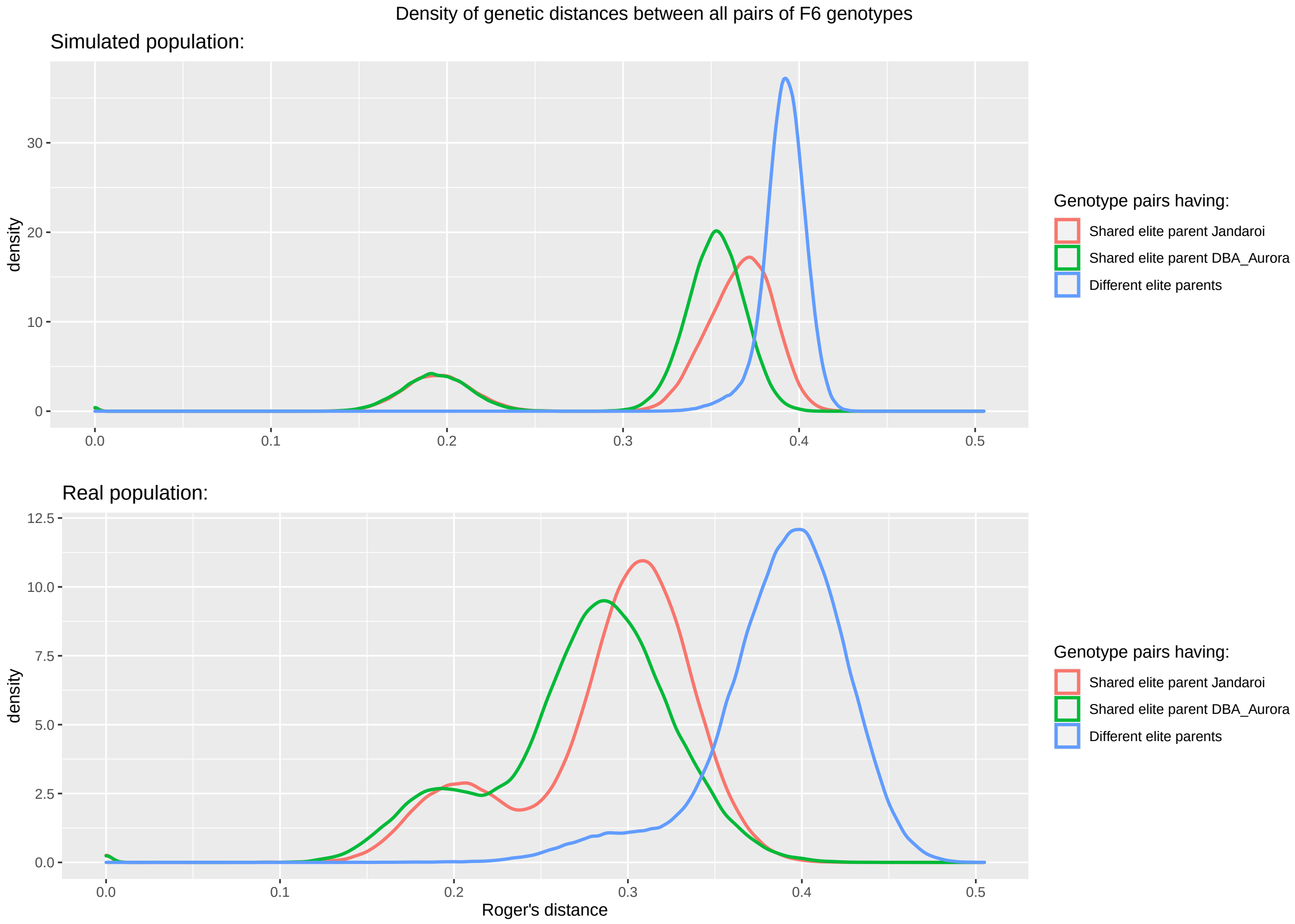


Figure S2: Distributions of genetic distance relationships between different structural elements of the simulated and empirical NAM populations. The genetic (Roger's) distance between each pair of final-generation genotypes was calculated, and each pair was either classified as: both genotypes being developed from elite parent Jandaroi, both genotypes being developed from elite parent DBA_Aurora, or the two genotypes being developed from different elite parents. The profiles of the distributions match across the simulated and real datasets, though the spread of the distance measures is greater in the real population.

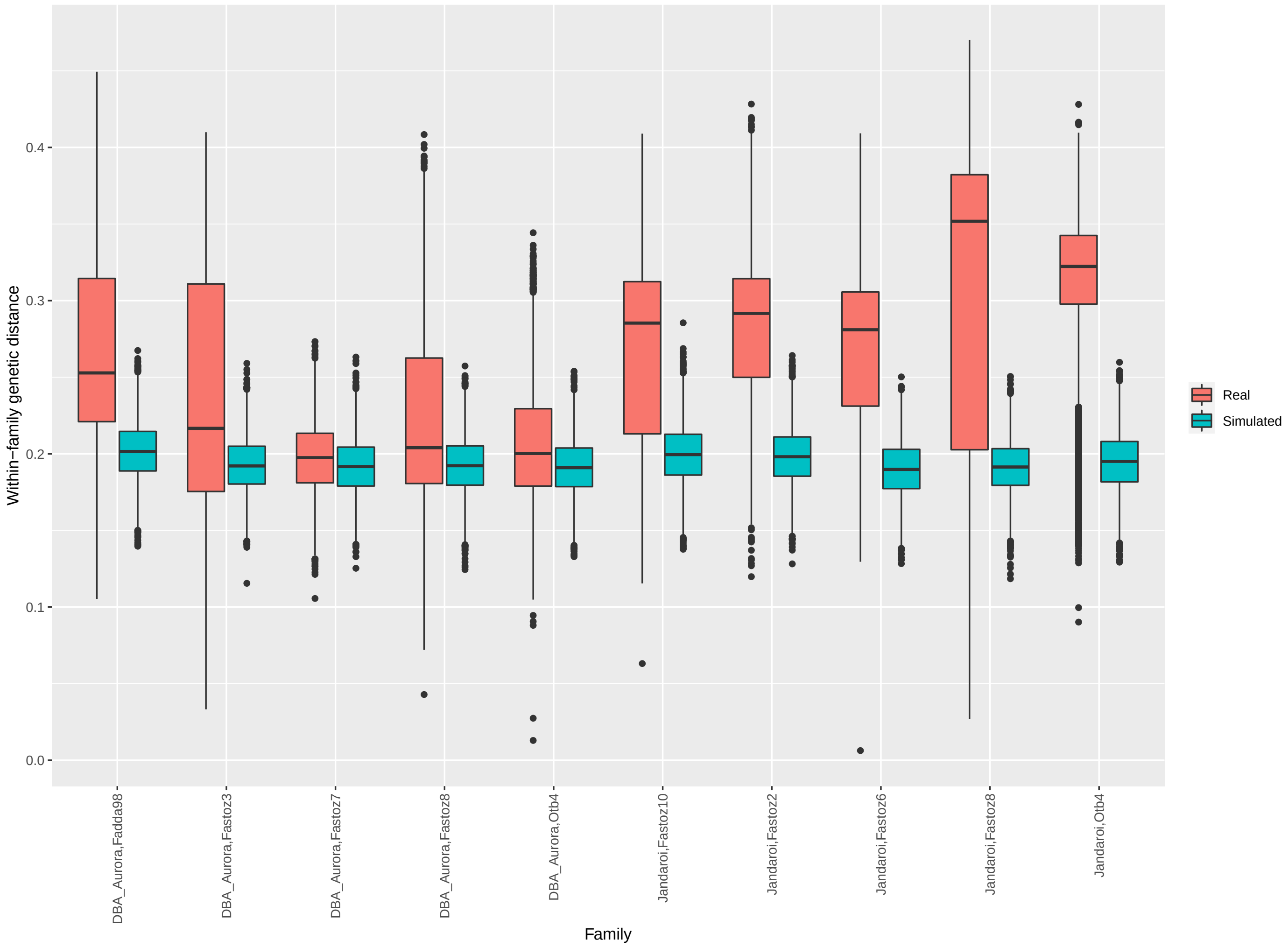


Figure S3: Diversity within families in simulated and empirical NAM program results. The genetic (Roger's) distance between each pair of final-generation genotypes was calculated, and boxplots plotted of the distribution of distance scores calculated between two members of the same family. The simulated dataset shows less within-family variation and a more consistent median within-family distance than the real dataset.
